# Metagenomics reveals fibre fermentation and AMR pathways in red grouse (*Lagopus scotica*) microbiota

**DOI:** 10.1101/2025.04.02.646739

**Authors:** Anum Ali Ahmad, Kathy Fletcher, Nicholas Hesford, Laura Glendinning

## Abstract

**Background:** The avian caecal microbiota plays a vital role in host nutrition, enabling non-digestible, fibrous material to be converted into compounds that can be absorbed and used as an energy source by the host. The diet of adult red grouse (*Lagopus scotica*) is dominated by heather (*Calluna vulgaris*), which is particularly high in fibre. It is therefore likely that the caecal microbiota plays a key role in enabling grouse to thrive on this diet. In this study, we present the first characterisation of the caecal microbiota of red grouse using modern sequencing methods.

**Results:** We performed metagenomic sequencing on caecal content samples from fifteen red grouse from three upland estates in Scotland. From this data, we constructed and characterised twelve high-quality, species-level metagenome assembled genomes (MAGs). Eleven of these MAGs could not be assigned a taxonomic label at species level, indicating that they may be novel species. MAGs belonged to diverse taxa (5 phyla) and several encoded genes and pathways for the digestion of fibres, including cellulose, hemi-cellulose, xylooligosaccharides and pectin. Several MAGs also contained antimicrobial resistance genes, predominantly related to vancomycin resistance.

**Conclusions:** This study is the first to reconstruct commensal microbial genomes from red grouse. The caeca contain diverse, often novel, microbial taxa capable of fermenting various fibres, potentially aiding in the digestion of the red grouse’s high-fibre diet. Further research is necessary to explore how these bacteria support red grouse nutrition and health.

## Background

The avian gut microbiota plays a vital role in maintaining host health and contributing to host nutrition (1). While the vast majority of avian microbiota studies have been conducted in farmed species, there are an increasing number of studies examining the microbiota of wild bird populations. These studies have shown that various environmental and genetic factors are important influencers of the composition and function of the gut microbiota (2).

An important challenge when studying wild species is acquiring samples that are truly representative of the gut microbiota. Studies that use wild birds are often limited to collecting non-invasive samples, such as faeces and caecal swabs. However, these samples do not quantitatively represent the microbiota of the large and small intestine (3, 4). Obtaining samples directly from the gastrointestinal section of interest is therefore advantageous, but is often not possible as it requires killing the bird. Game bird species offer a potential avenue for studying the microbiota in “wild” or minimally-managed avian populations, where gastrointestinal samples can be obtained in an ethical manner. Game birds are regularly shot during hunts, or to collect samples to monitor population health. The gastrointestinal tracts from these birds are usually discarded and can therefore be seen as a waste product of this industry. This allows researchers to access gastrointestinal samples without requiring additional animal sacrifice.

Diet has been reported as the most important factor affecting the wild avian microbiota (5). Most game bird species consume plants as a large component of their diet. For wild avian species that consume a predominantly or partially plant-based diet, it is likely that the gut microbiota plays an important role in enabling the host to extract energy from plant fibres. Members of the gut microbiota can ferment fibre into short chain fatty acids that can be used as an energy source by the host (6). In avians the vast majority of fibre fermentation occurs in the caeca (7), blind pouches located near the end of the gastrointestinal tract. In general, herbivorous avian species have longer caeca than non-herbivorous species (8).

Red grouse (*Lagopus scotica*) represent a particularly interesting game bird species in which to study microbiota fibre fermentation. The main component of the adult red grouse diet is heather (*Calluna vulgaris*), which is high in fibre (25%) and low in protein (7%) (9). Red grouse have long caeca (∼75 cm), which increase in size when they are supplied with a higher fibre diet (10). From feed digestibility studies, there is some evidence that the red grouse microbiota is able to digest a wide variety of fibres, including those that are generally considered more resistant to fermentation (e.g. lignin) (9).

There is also increasing interest in studying antimicrobial resistance in wild-living animal populations through a one health lens (11). As game birds are frequently consumed by humans as food, the presence of AMR in these birds is of interest from this perspective. Previous studies have identified AMR in gut-associated microbes in UK pheasants and partridges (12, 13), but this has not been examined in red grouse.

The gut microbiota has been studied in several grouse species using 16S rRNA gene sequencing and/or metagenomic sequencing, including in capercaillies (14, 15), greater sage grouse (16–18), wild prairie grouse (19) and ptarmigans (20–22). However, while bacteria have been cultured from the gastrointestinal tracts of red grouse (23, 24), their microbiota has not been characterised using these modern sequencing methods. In this study, we perform metagenomic sequencing analysis of red grouse caecal content samples. We construct twelve high-quality bacterial metagenome assembled genomes, and characterise their functional potential to metabolise fibre and their AMR potential.

## Methods

### Sample collection and processing

Samples were collected from red grouse killed during legal game bird shoots within the open season in three upland estates in Scotland. Samples were collected in the 2023 Scottish red grouse (*Lagopus scotica*) shooting season (August-October), at three distinct areas of moorland managed for recreational hunting of red grouse (Latitude and longitude: 55.75 - 3.01 (Samples RS1-5), 55.98 -3.33 (Samples RB1-5), and 57.02 -4.12 (Samples CA1-CA5)). Samples were collected from five birds per area. The intestines were removed from birds and frozen on site using a standard domestic freezer within 24 hours. Samples were then transported to the Roslin Institute within one month of sample collection, and stored at -80°C until DNA extraction. DNA was extracted from caecal contents using the QIAamp PowerFecal Pro DNA Kit according to the manufacturer’s instructions, using 250-300 mg of caecal contents. DNA quality and quantity were assessed using a NanoDrop spectrophotometer and a Qubit Fluorometer with a Qubit dsDNA Quantification Assay Kit (broad range). Shotgun metagenomic library preparation and sequencing was performed by Novogene, using an Illumina NovaSeq 6000 producing 150 bp paired-end reads. Samples RS1 and RS4 were produced on two separate sequencing runs in order to produce sufficient data quantity, then fastq files from the same sample were concatenated prior to analysis.

### Bioinformatic analysis

Sequences were checked for quality using FastQC (v.0.12.1) (25) and MultiQC (v.1.6) (26). Quality control was performed using fastp (v.0.24) (27), with the options “--detect_adapter_for_pe -l 30 -q 20 --trim_poly_g --trim_poly_x”. The host genome and feed genome were removed from adaptor trimmed read files. There is currently no RefSeq or GenBank genome available for *Lagopus scotica*, therefore the *Lagopus muta* (rock ptarmigan) genome was downloaded from RefSeq (GCF_023343835.1) and used instead of a host genome. Heather (*Calluna vulgari*) is the main food source for adult red grouse (28); therefore the *Calluna vulgari* genome was downloaded from GenBank (GCA_964145215.1). Sample reads were mapped to host and feed reference genomes using BWA-MEM (v.0.7.18) (29), followed by SAMtools (v.1.21, samtools fastq -f 12) (30) to select reads where both paired-end reads were unmapped. The unmapped reads were taxonomically classified using Kraken2 (v.2.3.1) (31) with a custom database built using *Lagopus muta* and bacterial, fungal, archaeal, viral, plant, plasmid, and protozoal RefSeq genomes (built on 23^rd^ June 2025). Bracken (v.3.0.1) (32) was then used to estimate their relative abundances (options: -k 35 -l 150). Single-sample assemblies and coassemblies were constructed using Megahit (v.1.2.9) (33).

CoverM (v.0.7) (34) was used to create depth files, which were used as input for genome binning by MetaBAT 2 (v.2.17) (35). Contamination and completeness of bins were estimated using CheckM2 (v.1.0.2) (36). dRep (v.3.5.0) (37) was used to dereplicate genome bins into MAGs, with minimum completeness set at 80% and maximum contamination at 10%. Genomes were dereplicated at 95% ANI for species-level MAGs, and 99% ANI for strain-level MAGs. Viruses were identified from megahit assemblies using VirSorter2 (v.2.2.4: database downloaded 3^rd^ February 2025) (38) (options: --include-groups dsDNAphage,ssDNA --min-length 5000 --min-score 0.5 -j 28 all).

GTDB-Tk (v.2.4.0) (39) was used to assign taxonomy to MAGs (database downloaded 24^th^ January 2025). METABOLIC was used to assign metabolic function to MAGs (v.4.0) (40). Antimicrobial resistance (AMR) genes present in MAGs and megahit assemblies were identified using RGI main (v. 6.0.3) (41) with the Comprehensive Antibiotic Resistance Database (database downloaded 17^th^ March 2025). Loose hits were included, and the -- low_quality option was used for megahit assemblies. The abundance of MAGs within samples was calculated using CoverM (options: -p bwa-mem -m mean relative_abundance trimmed_mean covered_bases variance length reads_per_base count rpkm tpm --min-read-aligned-percent 75 --min-read-percent-identity 95 --min-covered-fraction 0). Graphs and data summaries were produced in R using the packages reshape2 (v.1.4.4) (42), cowplot (v.1.1.3) (43), dplyr (v.1.1.4) (44), ggplot2 (v.3.5.1) (45) and tidyverse (v.2.0.0) (46). MAG taxonomic trees were constructed using PhyloPhlAn (v.3.1.1) (47) (options: --diversity high -f supermatrix_aa.cfg --subsample phylophlan –fast). Taxonomic trees were midpoint rerooted and visualised using figtree (v.1.4.3) (48).

## Results

### Sequence quality

We performed metagenomic sequencing on caecal content samples from fifteen Scottish red grouse. After removal of poor-quality sequences, and trimming of adaptors, polyAs and polyGs, there were on average 102,686,250 ± 24,122,717 reads per sample. After removal of host and feed sequences the average number of reads per sample dropped by 89%, to 10,902,109 ± 7,733,350. This was predominantly due to the removal of host reads, indicating that our samples had a large proportion of host contamination. From single sample assemblies and a coassembly, we constructed 161 putative genome bins. Quality-filtering and dereplication resulted in twelve high-quality (<10% contamination, >80% complete) species-level MAGs (**Table 1**). We also constructed strain-level MAGs, but as we found only one more MAG than at species-level we decided to restrict our further analyses to species level. Eighteen high quality viral genomes were identified (**Table S1**), two of which were proviruses.

**Table 1:**
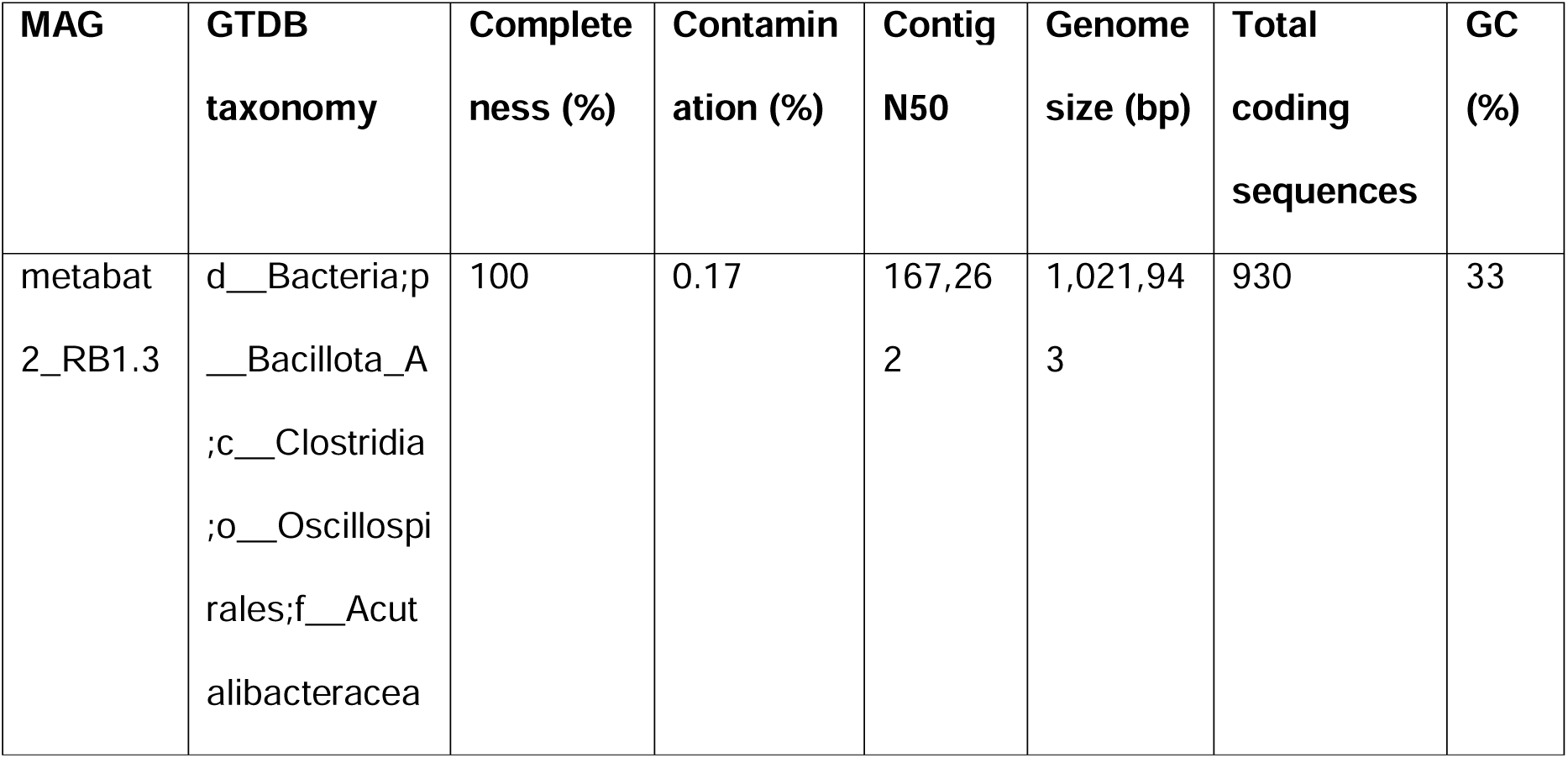
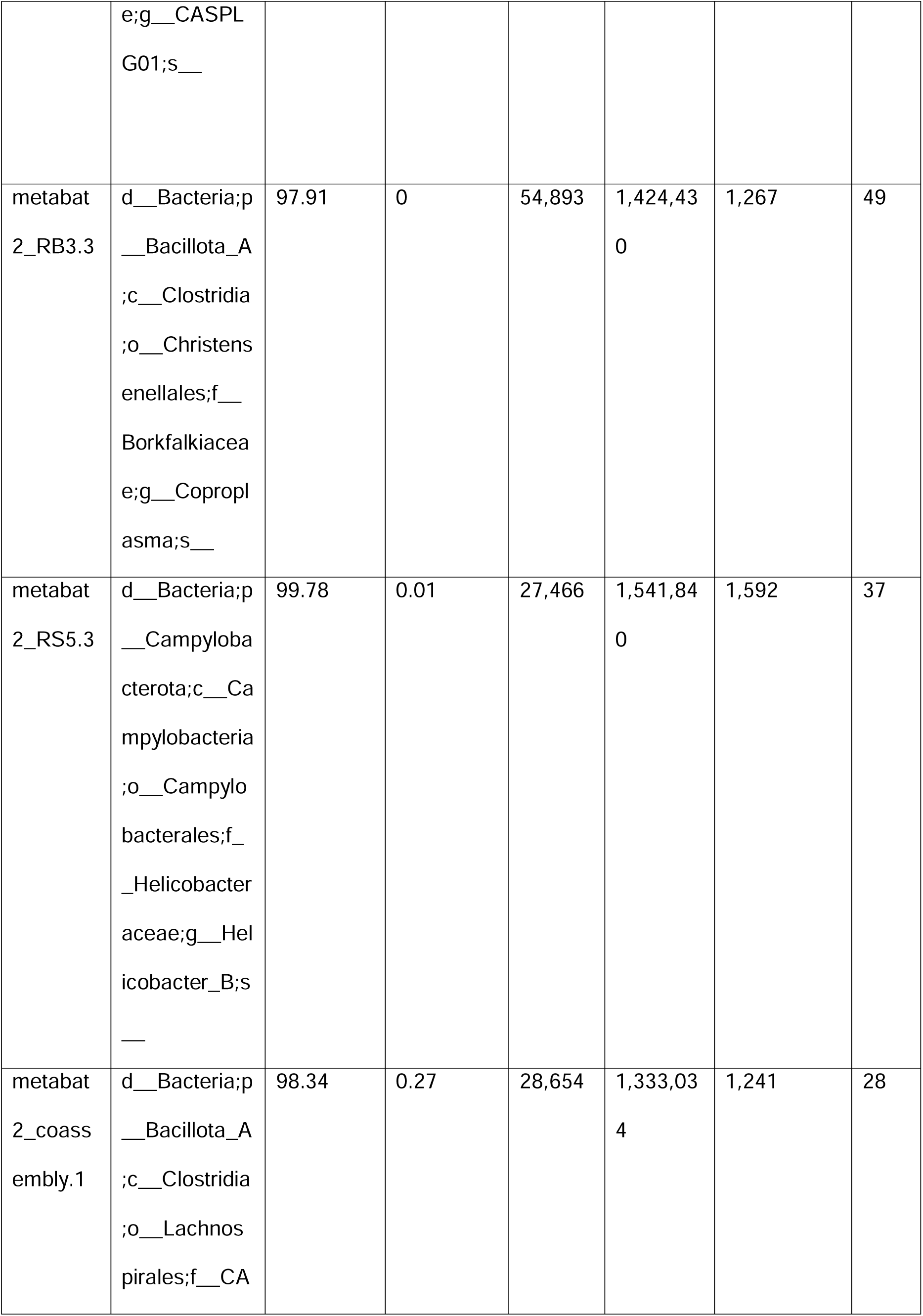
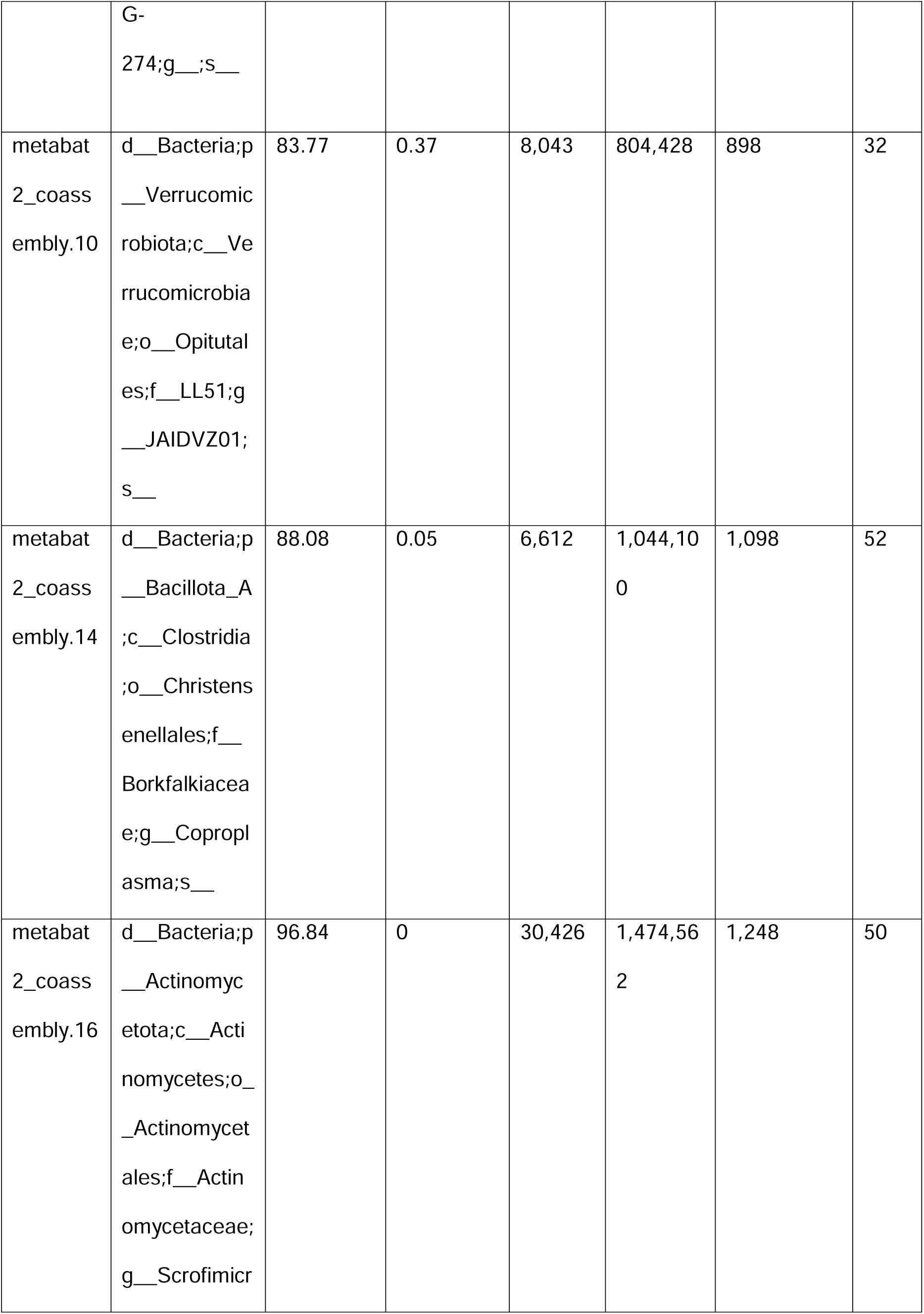
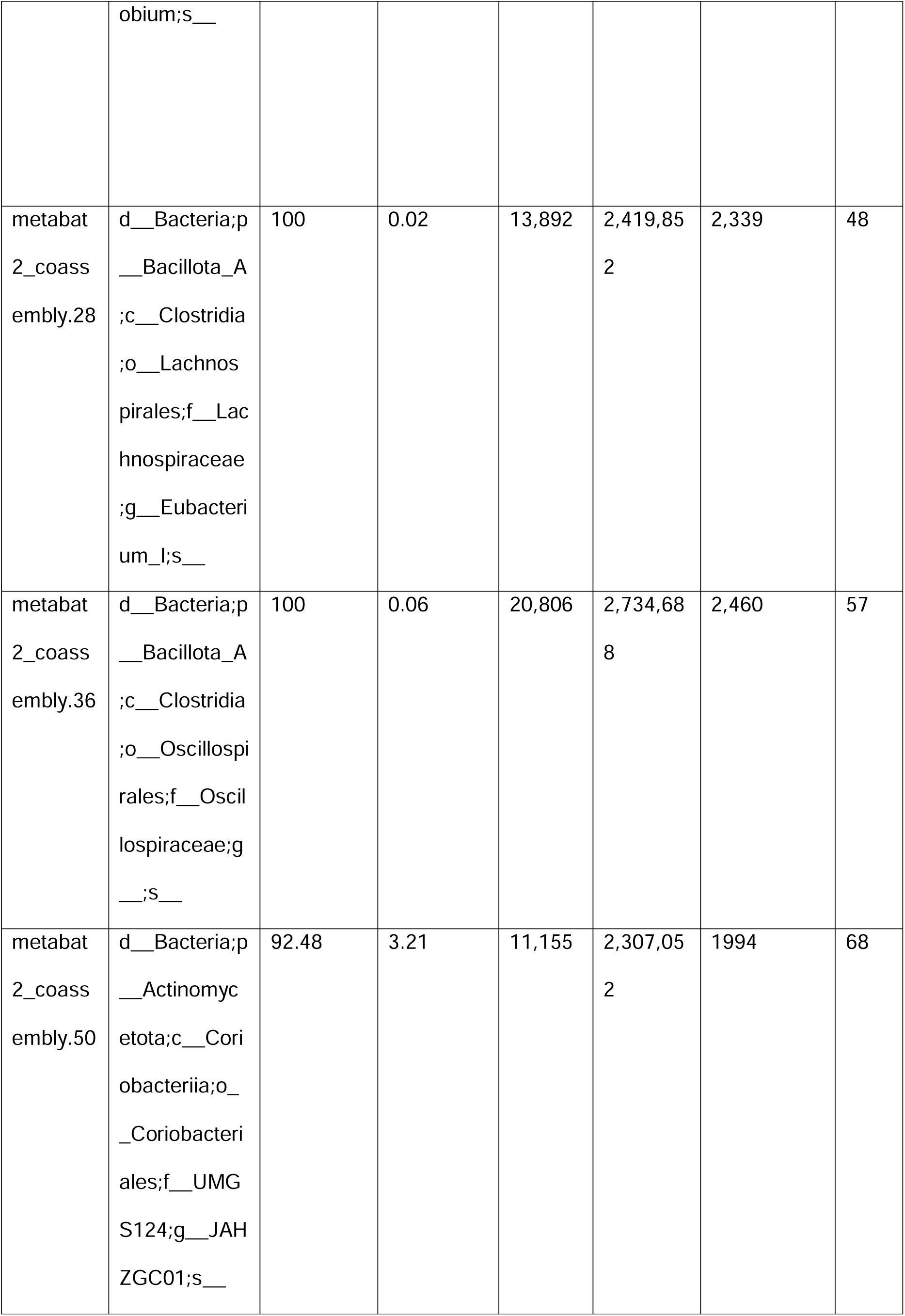
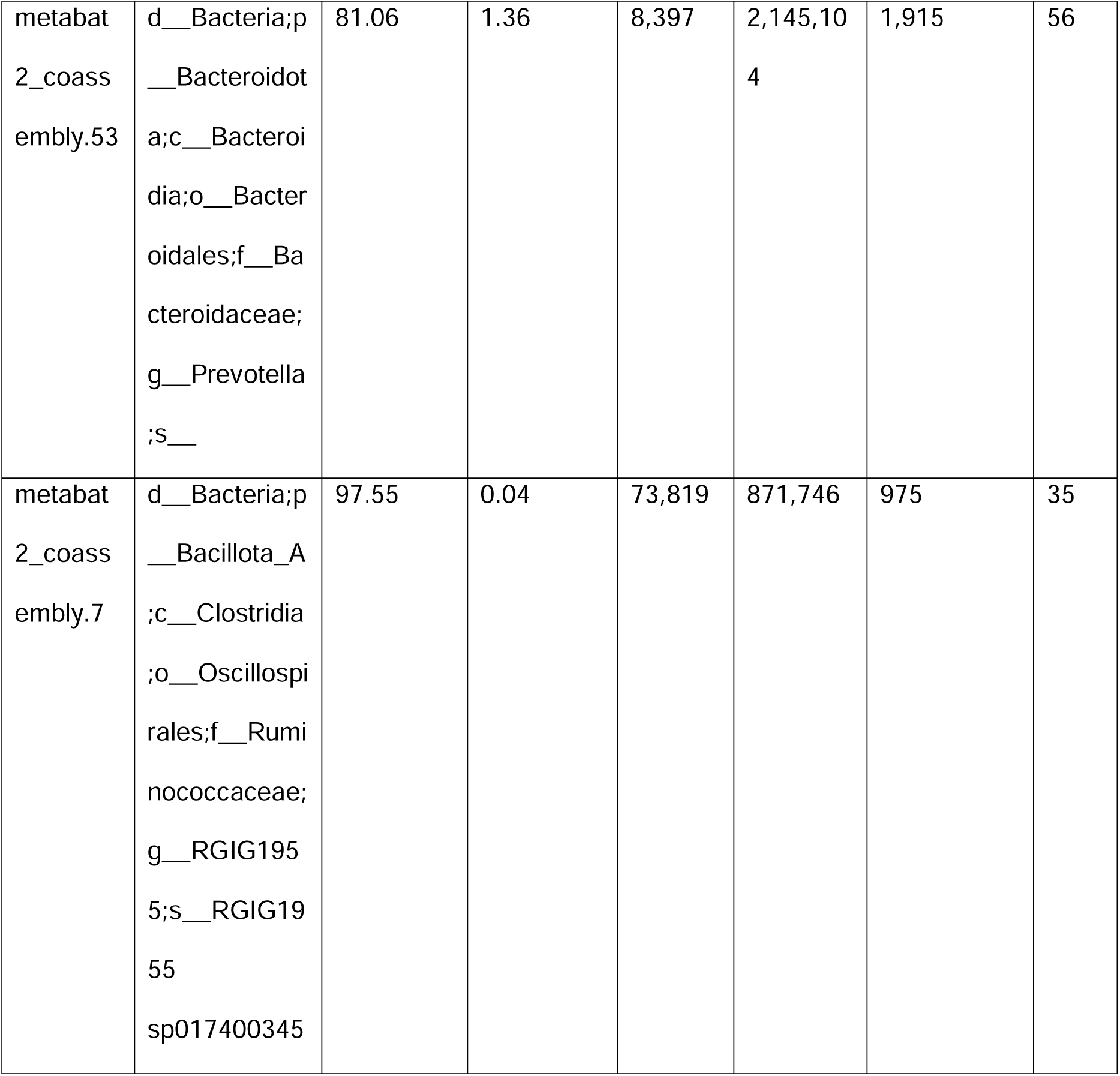
Genome stats for species-level MAGs.

### Taxonomy of MAGs

All MAGs belonged to bacteria (**Figure 1**), with the most common phylum being Bacillota_A (n=7), all of the members of which belonged to the class Clostridia. Two members of this phylum were identified as members of the genus *Coproplasm*a, within the family Christensenellales. The remaining MAGs in this phylum belonged to Lachnospirales (one identified to family CAG-274 and one identified to the genus *Eubacterium*), and Oscillospirales (one identified to family Oscillospiraceae, one identified to genus *CASPLG01*, and one identified as *RGIG1955 sp017400345*). This latter MAG was the only one from our dataset to be identified to species level by GTDB-Tk. Two MAGs were identified as members of the phylum Actinomycetota, and were assigned to the genera *Scrofimicrobium* and *JAHZGC01*. One MAG belonged to the phylum Bacteroidota (genus *Prevotella*), one belonged to the phylum Campylobacterota (genus *Helicobacter_B*) and one belonged to the phylum Verrucomicrobiota (genus *JAIDVZ01*).

**Figure 1:**
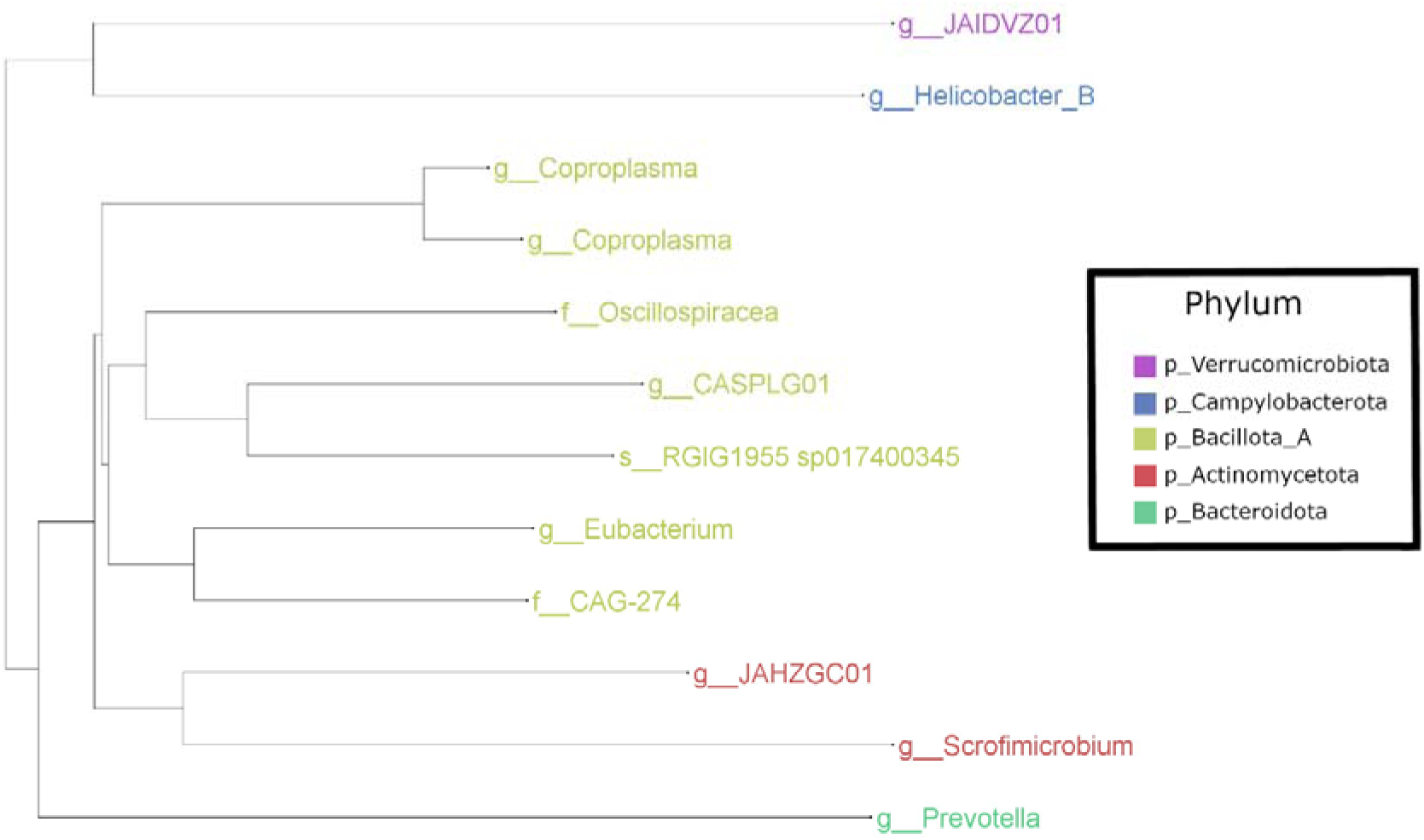
Taxonomic tree of species-level, high-quality metagenome assembled genomes constructed from red grouse caecal metagenomic data. Tip labels represent the lowest taxonomy assigned to each MAG by GTDB-Tk.

For most samples, the majority of reads did not map to MAGs (69.5 ± 14.3% relative abundance) (**Figure 2**). By far the most abundant MAG was metabat2_RS5.3 (genus *Helicobacter_B*) (26.3 ± 15.4%), followed by metabat2_RB1.3 (genus *CASPLG01*) (1.9 ± 1.3%) and metabat2_RB3.3 (genus *Coproplasma*) (1.4 ± 1.9%). Some samples contained particularly high levels of *Helicobacter_B*, including RB2 (64.5%), RB4 (46.7%) and RB5 (44.0%).

**Figure 2:**
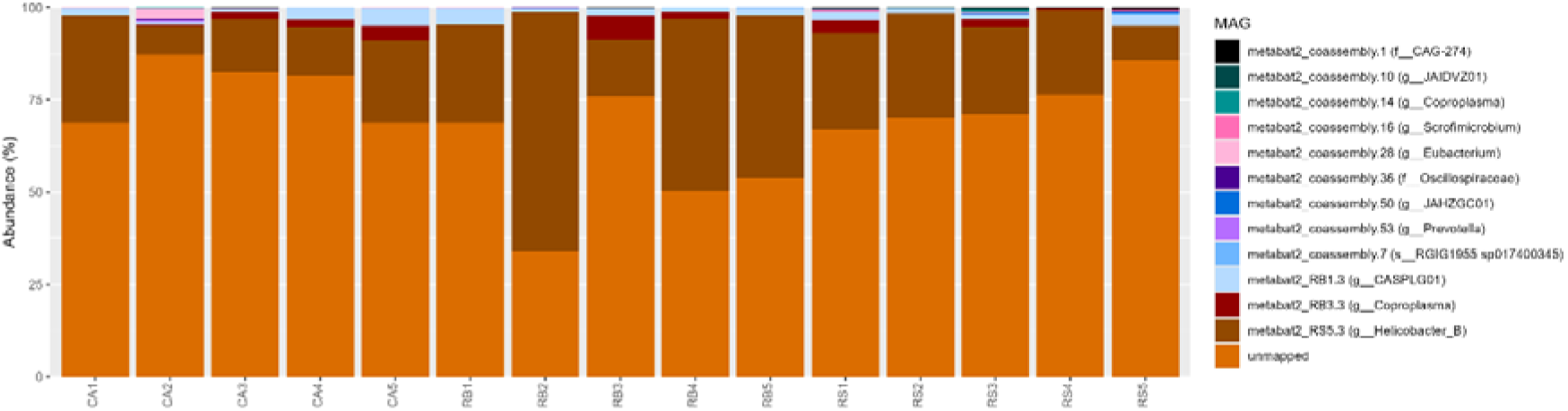
Relative abundance of species-level, high-quality metagenome assembled genomes in red grouse caecal samples. Taxonomies represent the lowest taxonomy assigned to each MAG by GTDB-Tk.

### Metabolism of MAGs

Carbohydrate active enzymes (CAZymes) play a key role in carbohydrate fermentation by the microbiota, including the fermentation of fibrous compounds. Overall, 40 different CAZymes were identified in our MAGs. The most common CAZyme (18 genes) was GH13, followed by GH3 (16 genes), GH25 (11 genes), GH2 (9 genes), GH43 (8 genes), GH18, (7 genes), GH77 (7 genes), GH23 (5 genes), GH28 (5 genes). All other CAZymes had less than 5 genes present in our MAGs (**Table S2**). MAG metabat2_coassembly.53 (*Prevotella*) contained substantially more CAZyme genes than any other MAG: 55 genes belonging to 27 CAZyme families. Metabat2_coassembly.28 (*Eubacterium*) and metabat2_coassembly.36 (Oscillospiraceae) also contained a large number of CAZyme genes: 25 genes from 17 CAZyme families and 20 genes from 13 families respectively. The three most abundant MAGs in our samples contained far fewer CAZyme genes: metabat2_RS5.3 (*Helicobacter_B*) contained 3 genes from 2 families, metabat2_RB1.3 (*CASPLG01*) contained 7 genes from 4 families and metabat2_RB3.3 (*Coproplasma*) contained 9 genes from 5 families.

Several MAGs were predicted to be able to metabolise complex carbohydrates commonly found in plants. Cellulose degradation via beta-glucosidase was present in five of our MAGs, including the taxa *Coproplasma*, *Eubacterium*, Oscillospiracea and *Prevotella*. Cellulose degradation via cellobiosidase or cellulase was not present in our dataset. Hemicellulose-debranching enzymes were also present amongst our MAGs, including arabinosidase (*Prevotella*), beta-glucuronidase (*Coproplasma* and *JAHZGC01*) and alpha-L-rhamnosidase (Oscillospiraceae and *Prevotella*). One endohemicellulase gene (alpha-D-xyloside xylohydrolase) was also identified in our *Prevotella* MAG. Beta-xylosidase (targeting xylooligosaccharides) was also only identified in this *Prevotella* MAG.

Heather is known to contain various phenolic compounds. Enzymes for converting phenol into benzoyl-CoA were found in two MAGs (Oscillospiraceae and *Helicobacter_B*). Although starch is present in low concentrations in heather, amylolytic enzymes involved in starch digestion were also present in two MAGs: alpha-amylase in the *Prevotella* MAG and pullulanase in the *JAIDVZ01* MAG. Only two genes were identified related to sulphur metabolism, namely sulphite reduction in the *Eubacterium* MAG, and sulphur oxidation in the *CAG-274*.

### Antimicrobial resistance genes

We next investigated whether our MAGs contained antimicrobial resistance (AMR) genes (**Table S3**). Whilst we do not expect any of the sampled red grouse to have been directly treated with antibiotics, AMR genes occur naturally in many bacteria, even without antibiotic exposure. The RGI “strict” algorithm is used to detect previously unknown variants of known AMR genes. Twenty-nine AMR genes were detected in our MAGs using this algorithm (**Table 2**), the vast majority of which encoded resistance to vancomycin (27 genes in 9 MAGs). A metronidazole resistance gene was detected in metabat2_RB3.3 (*Coproplasma*) and a tetracycline gene in metabat2_RS5.3 (*Helicobacter*).

**Table 2:** AMR genes present in MAGs (strict RGI hits)

| MAG | Taxonomy | Antibiotic | Gene | % Identity | Resistance Mechanism | AMR Gene Family |
| --- | --- | --- | --- | --- | --- | --- |
| metabat2_coassembly.1 | f__CAG-274 | vancomycin;<br>teicoplanin | vanW | 30.04 | antibiotic target alteration | vanW;<br>glycopeptide resistance gene cluster |
| metabat2_coassembly.1 | f__CAG-274 | vancomycin | vanG | 41.88 | antibiotic target alteration | glycopeptide resistance gene cluster;<br>Van ligase |
| metabat2_<br>coassembl<br>y.1 | f__CAG-274 | vancomyci<br>n | vanT | 36.04 | antibiotic<br>target<br>alteration | glycopepti<br>de<br>resistance<br>gene<br>cluster;<br>vanT |
| metabat2_<br>coassembl<br>y.14 | g__Copro<br>plasma | vancomyci<br>n;<br>teicoplanin | vanW | 30.75 | antibiotic<br>target<br>alteration | vanW;<br>glycopepti<br>de<br>resistance<br>gene<br>cluster |
| metabat2_<br>coassembl<br>y.28 | g__Eubact<br>erium | vancomyci<br>n;<br>teicoplanin | vanW | 32.82 | antibiotic<br>target<br>alteration | vanW;<br>glycopepti<br>de<br>resistance<br>gene<br>cluster |
| metabat2_<br>coassembl<br>y.28 | g__Eubact<br>erium | vancomyci<br>n | vanT | 38.24 | antibiotic<br>target<br>alteration | glycopepti<br>de<br>resistance<br>gene<br>cluster;<br>vanT |
| metabat2_<br>coassembl<br>y.36 | f__Oscillo<br>spiraceae | vancomyci<br>n;<br>teicoplanin | vanW | 39.9 | antibiotic<br>target<br>alteration | vanW;<br>glycopepti<br>de |
|  |  |  |  |  |  | resistance<br>gene<br>cluster |
| metabat2_<br>coassembl<br>y.36 | f__Oscillo<br>spiraceae | vancomyci<br>n | vanT | 35.22 | antibiotic<br>target<br>alteration | glycopepti<br>de<br>resistance<br>gene<br>cluster;<br>vanT |
| metabat2_<br>coassembl<br>y.36 | f__Oscillo<br>spiraceae | vancomyci<br>n | vanY | 61.62 | antibiotic<br>target<br>alteration | vanY;<br>glycopepti<br>de<br>resistance<br>gene<br>cluster |
| metabat2_<br>coassembl<br>y.36 | f__Oscillo<br>spiraceae | vancomyci<br>n;<br>teicoplanin | vanW | 39.59 | antibiotic<br>target<br>alteration | vanW;<br>glycopepti<br>de<br>resistance<br>gene<br>cluster |
| metabat2_<br>coassembl<br>y.36 | f__Oscillo<br>spiraceae | vancomyci<br>n | vanG | 39.94 | antibiotic<br>target<br>alteration | glycopepti<br>de<br>resistance<br>gene<br>cluster;<br>Van ligase |
| metabat2_<br>coassembly.36 | f__Oscillo<br>spiraceae | vancomycin | vanT | 48.37 | antibiotic<br>target<br>alteration | glycopeptide<br>resistance<br>gene<br>cluster;<br>vanT |
| metabat2_<br>coassembly.36 | f__Oscillo<br>spiraceae | vancomycin | vanG | 53.78 | antibiotic<br>target<br>alteration | glycopeptide<br>resistance<br>gene<br>cluster;<br>Van ligase |
| metabat2_<br>coassembly.36 | f__Oscillo<br>spiraceae | vancomycin | vanW | 56.25 | antibiotic<br>target<br>alteration | vanW;<br>glycopeptide<br>resistance<br>gene<br>cluster |
| metabat2_<br>coassembly.50 | g__JAHZ<br>GC01 | vancomycin | vanXY | 41.85 | antibiotic<br>target<br>alteration | glycopeptide<br>resistance<br>gene<br>cluster;<br>vanXY |
| metabat2_<br>coassembly.50 | g__JAHZ<br>GC01 | vancomycin | vanG | 40.51 | antibiotic<br>target<br>alteration | glycopeptide<br>resistance gene<br>cluster;<br>Van ligase |
| metabat2_<br>coassembly.50 | g__JAHZ<br>GC01 | vancomycin | vanT | 40.51 | antibiotic<br>target<br>alteration | glycopeptide<br>resistance<br>gene<br>cluster;<br>vanT |
| metabat2_<br>coassembly.50 | g__JAHZ<br>GC01 | metronidazole | nimJ | 49.07 | antibiotic<br>inactivation | nitroimidazole<br>reductase |
| metabat2_<br>coassembly.50 | g__JAHZ<br>GC01 | vancomycin | vanH | 43.71 | antibiotic<br>target<br>alteration | vanH;<br>glycopeptide<br>resistance<br>gene<br>cluster |
| metabat2_<br>coassembly.53 | g__Prevotella | vancomycin | vanT | 36.98 | antibiotic<br>target<br>alteration | glycopeptide<br>resistance<br>gene<br>cluster;<br>vanT |
| metabat2_<br>coassembly.7 | s__RGIG1<br>955<br>sp017400 | vancomycin | vanY | 37.74 | antibiotic<br>target<br>alteration | vanY;<br>glycopeptide |
|  | 345 |  |  |  |  | resistance<br>gene<br>cluster |
| metabat2_<br>coassembly.7 | s__RGIG1<br>955<br>sp017400<br>345 | vancomycin | vanG | 61.21 | antibiotic<br>target<br>alteration | glycopeptide<br>resistance<br>gene<br>cluster;<br>Van ligase |
| metabat2_<br>coassembly.7 | s__RGIG1<br>955<br>sp017400<br>345 | vancomycin | vanXY | 53.66 | antibiotic<br>target<br>alteration | glycopeptide<br>resistance<br>gene<br>cluster;<br>vanXY |
| metabat2_<br>coassembly.7 | s__RGIG1<br>955<br>sp017400<br>345 | vancomycin | vanT | 53.12 | antibiotic<br>target<br>alteration | glycopeptide<br>resistance<br>gene<br>cluster;<br>vanT |
| metabat2_<br>RB1.3 | g__CASP<br>LG01 | vancomycin | vanT | 37.67 | antibiotic<br>target<br>alteration | glycopeptide<br>resistance<br>gene<br>cluster;<br>vanT |
| metabat2_<br>RB1.3 | g__CASP<br>LG01 | vancomycin | vanG | 41.13 | antibiotic<br>target<br>alteration | glycopeptide<br>resistance<br>gene<br>cluster;<br>Van ligase |
| metabat2_<br>RB1.3 | g__CASP<br>LG01 | vancomycin | vanY | 36.21 | antibiotic<br>target<br>alteration | vanY;<br>glycopeptide<br>resistance<br>gene<br>cluster |
| metabat2_<br>RB3.3 | g__Copro<br>plasma | vancomycin;<br>teicoplanin | vanW | 28.37 | antibiotic<br>target<br>alteration | vanW;<br>glycopeptide<br>resistance<br>gene<br>cluster |
| metabat2_<br>RS5.3 | g__Helico<br>bacter | Tetracycline | adeF | 43.68 | antibiotic<br>efflux | resistance<br>-<br>nodulation<br>-cell<br>division<br>(RND)<br>antibiotic<br>efflux<br>pump |

The RGI “loose” algorithm aims to identify more distant homologs of AMR genes. While this method is more prone to false positives, we decided to cautiously employ it as members of the grouse microbiota are likely to be poorly represented in the CARD database. This method identified 13,159 potential AMR gene homologs amongst our MAGs, with the top three targeted antibiotics as oxacillin, tetracycline and vancomycin. The MAGs with the most putative AMR gene homologs were metabat2_coassembly.36 (n=1972, Oscillospiraceae), metabat2_coassembly.28 (n=1882, *Eubacterium*), and metabat2_coassembly.50 (n=1525, *JAHZGC01*).

### Taxonomic classification of reads

As MAGs normally only capture the most abundant bacterial and archaeal taxa in samples, we used Kraken2/Bracken to taxonomically classify our metagenomic reads and characterise taxonomies that may not have been captured by MAGs (**Figure 3**). On average, 85.0 ± 1.63% metagenomic reads remained unclassified in our data. The classified reads were assigned to 4 domains, 82 phyla, and 2,632 genera across all samples. At the domain level, the reads were dominated by bacteria (56.3 ± 12.8%, relative abundance) and Eukaryota (43.2 ± 12.7%) across all samples **(Figure 3A)**. Streptophyta (40.3 ± 12.0%) and Campylobacterota (27.7 ± 13.4 %) were dominant phyla **(Figure 3B)**. Other minor phyla included Pseudomonadota (9.27 ± 3.01%), Bacillota (8.87 ± 3.25%), Bacteroidota (4.41 ± 2.84%), and Actinomycetota (3.49 ± 2.87%). At the genus level, *Helicobacter* (18.8 ± 9.42%) and *Campylobacter* (6.21 ± 2.89%) were highly abundant across all samples **(Figure 3C)**. Other minor genera included *Cryptomeria* (2.28 ± 0.57%), *Escherichia* (2.18 ± 2.35%), *Solanum* (1.88 ± 0.69%), and *Cucumis* (1.14 ± 0.52%).

**Figure 3:**
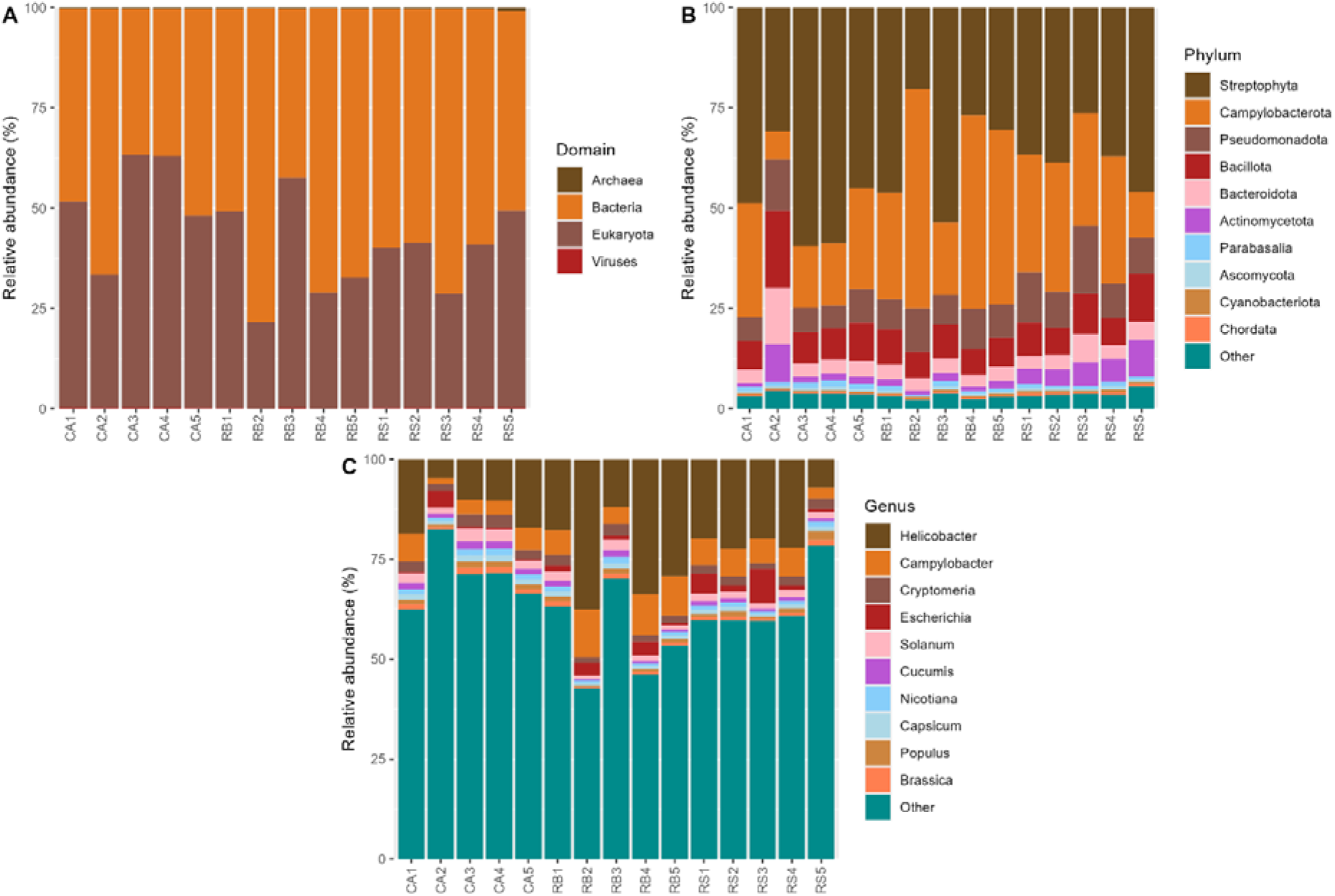
Barplot showing Kracken2/Bracken classified metagenomic reads at the domain (A), phylum (B), and genus (C) levels. The top 10 highly abundant taxa are displayed, while the remaining taxa are grouped as “Other”.

## Discussion

The red grouse represents a particularly interesting species in which to study the microbiota, due to its reliance on a single, high-fibre plant species (heather - *Calluna vulgaris*) for the majority of its diet (9). Avians do not produce fibre fermenting enzymes, and therefore in order to acquire nutrition from fibre they are reliant on their gut microbiota to ferment fibre into compounds that they can absorb and use as an energy source (e.g. short chain fatty acids) (6). In this study, we examined the gut microbiota of red grouse for the first time using metagenomics to construct metagenome assembled genomes. Unlike metabarcoding methods (e.g. 16S rRNA gene sequencing), metagenomics allows us to not only study the taxa that are present in a community, but also provides functional information on their metabolic and virulence potential.

Taxonomic profiling using Kraken2/Bracken revealed that the majority of our metagenomic reads remained unclassified. This is likely due to the fact that the gut microbiota of red grouse has not previously been examined using metagenomic sequencing methods, so red grouse-specific members of the gut microbiota will not currently be included in genomic databases. Similar results have been reported for other wild bird species (49–51), demonstrating the understudied nature of the microbiota of these species. A large proportion of our reads originated from plants, likely from the bird’s diet. Due to our quality control methods these reads are unlikely to have impacted our microbiota analysis, but these results highlight the possibility of using gut metagenomics to study the diets of wild birds. The results from our Kraken2/Bracken analysis should be interpreted cautiously. As members of the red grouse microbiota are unlikely to be represented in the RefSeq database, they are unlikely to be present in the Kraken2 database used in our analysis. Misassignment of taxonomies is therefore likely, particularly at lower taxonomic levels.

We constructed MAGs to gain a deeper understanding of the microbial community composition and its functional potential, and to begin to address the lack of red grouse microbiota representatives in public sequence databases. In this study, we identified twelve high-quality, species-level MAGs. Of our twelve MAGs, eleven could not be assigned to a species by GDTB-Tk. The GTDB is constructed from high-quality genomes obtained from RefSeq and GenBank, and the release used in this study contains 596,859 microbial genomes in 113,104 species clusters, including genomes constructed from metagenomic data. If genomes cannot be assigned a taxonomy by this tool it is therefore a good indication that they are likely to be novel. Only one MAG was assigned a species label, RGIG1955 sp017400345 within the Ruminococcaceae family. This is a poorly characterised species, which has previously only been identified in metagenomics data from ruminants (52, 53). A MAG from the genus *Helicobacter_B* was the most abundant MAG in our samples, comprising on average 26% of the microbiota. Kraken2/Bracken analysis also identified this genus to be highly abundant in our metagenomic reads. *Helicobacter* is a highly diverse genus, which while commonly associated with gastrointestinal disease (e.g. *H.pylori* infection in humans) is also a common member of the gut microbiota in many species, including grouse (16, 21). In ptarmigan intestines, Bjørnsen *et al.* found similar average abundances of *Helicobacter* as observed in our study (24% ± 3) (54).

The majority of our MAGs were identified as Clostridia, which is a commonly identified taxon in the guts of other grouse species (16, 17, 20). Of these MAGs, most are from poorly described taxa identified from metagenomic data: the genus *Coproplasma* has been identified in the chicken gut (55), the genus *CASPLG01* has been identified in mice (56–58), and the family CAG-274 has been identified in metagenomic datasets from diverse species (59–61). In contrast, the diverse genus *Eubacterium* is relatively well studied, and has been proposed as a potential probiotic in humans due to the ability of some species to produce butyrate (62). Of the two MAGs assigned to the phylum Actinomycetota, one belonged to the genus *JAHZGC01* which has only one representative in the GTDB, obtained from human metagenomic data (63). The other MAG belonged to the genus *Scrofimicrobium*, which has been isolated from the gastrointestinal tracts of diverse species (64, 65).Only one MAG was identified as belonging to the phylum Verrucomicrobiota, a member of the genus *JAIDVZ01*, which is poorly characterised and has only been identified in metagenomic data (66, 67). Conversely, the only MAG identified in the phylum Bacteroidota belonged to the very well characterised genus *Prevotella*. This genus is known to play a key role in the fermentation of fibre in various herbivores (68) and is commonly found in the microbiota of avians, including grouse species (16, 17).

It is therefore unsurprising that our *Prevotella* MAG contained the largest quantity of CAZyme genes amongst our MAGs, and was predicted to degrade fibrous compounds including cellulose, hemi-celluloses and xylooligosaccharides. However, several MAGs from less well characterised taxa were also found to encode for diverse CAZyme genes and to be capable of degrading fibres, highlighting the potential role of poorly characterised taxa in fibre fermentation in the grouse caeca. Several CAZymes families with xylooligosaccharide, pectin and hemicellulose degrading activity (e.g. GH36, GH42, GH73) were identified both in our dataset and in the microbiota of other grouse species (14, 20). Various phenolic compounds are also found in heather, and have been shown to affect fermentation in the rumen microbiota (69). Pathways for the degradation of phenols were found in both our MAGs and other grouse species (17, 20), as were pathways for starch degradation (20). These findings indicate some degree of shared function in the grouse gut microbiota, in relation to digestion of plant-derived components of the diet.

Our MAGs also contained several antimicrobial resistance genes. It is likely that many of the hits found using the “loose” algorithm from RGI are false-positives, due to their high number. Using the strict RGI algorithm, the most commonly targeted antibiotic by AMR genes was vancomycin. This included the gene families: vanW, Van ligase, vanT, vanY, vanXY, and vanH. These genes are present in a wide range of taxa bacterial taxa (70) and have also been identified in samples from wild western capercaillies (14), but not amongst wild prairie grouse (19). Many of these resistance genes are not well characterised, with limited information on their molecular structures (70). vanT is found within resistance cassettes and encodes a membrane-bound serine racemase, while vanH encodes a dehydrogenase that converts pyruvate to d-lactate. The mechanism of action of vanW is currently unknown. vanY and vanXY are not required for vancomycin resistance but may contribute to increased resistance. A resistance-nodulation-cell division antibiotic efflux pump, predicted to target tetracycline was found in our *Helicobacter* MAG. Similar efflux pumps have been found in other species of *Helicobacter* (71). A nitroimidazole reductase gene was also identified in our *JAHZGC01* MAG. Similar genes can be found in a wide variety of bacterial taxa (72), but to our knowledge this is the first time this gene has been identified in this taxon. The ecological and one health relevance of these AMR genes is currently unknown, and is worth further investigation.

The main weakness of our study is the relatively small number of MAGs which we were able to construct in comparison to studies in other wild avians (14, 51, 73). However, it is worth noting that the MAG quality cut-offs used for these studies were often substantially lower than in our study (e.g. minimum 50% completeness). We chose to use a higher cut-off of 80% completeness as we were predominantly interested in predicting the metabolism of these bacterial species, and lower genome completeness can have a significant effect on functional inference (74). The small number of taxa observed in our samples may be truly representative of the microbiota, with the community being dominated by a small number of more abundant bacteria. In general, the more abundant a species is in a sample then the more likely it will be able to be constructed into a high-quality MAG. The high abundance of *Helicobacter* in our samples may have led to other species not being present in a high enough relative abundance to be constructed into MAGs. The low number of MAGs may also be due to reduced sequencing depth, due to the large amount of reads that belonged to host (89%).This percentage is similar to that found in saliva, throat, and vaginal swabs (75), but is unusual for the caeca or large intestine. One possible methodological explanation for this high degree of contamination could be how samples were stored. In order to enhance ease of sampling in the field and prevent spoiling of the carcass, intestines were initially stored whole, then defrosted and the contents removed for DNA extraction. This may have led to increased lysis of host cells, and thereby increased host DNA in our samples.

## Conclusion

This is the first study to reconstruct microbial genomes from the caeca of red grouse. The red grouse caeca harbours diverse microbial taxa, many of which are novel. These taxa have the potential to ferment various forms of fibre, thereby contributing to the digestion of the red grouse’s high fibre diet. Further studies are needed to understand how these bacteria contribute to grouse nutrition and health.

## List of abbreviations

AMR: Antimicrobial resistance
ANI: Average nucleotide identity
Bp: Base pairs
CAZymes: Carbohydrate active enzymes
GH: Glycoside hydrolases
GTDB: Genome taxonomy database
MAG: Metagenome assembled genome
rRNA: ribosomal RNA

## Declarations

### Ethics approval and consent to participate

Red grouse carcasses were privately owned by the estates on which the birds were shot. The Game Wildlife and Conservation Trust accessed and collected the gastrointestinal samples for this study. Informed consent was obtained from the owners to use these samples in our study. Field studies were conducted in accordance with local legislation. In the United Kingdom the birds listed in Schedule 2 of the Wildlife and Countryside Act 1981 (which include red grouse) are exempt from the general provision of protection. Any person is able to kill or ‘take’ a species listed outside of the close season for that bird. Therefore, red grouse can be legally taken between 12th August and 10th December within the United Kingdom. Red grouse were not killed for the purpose of providing samples for this study. Samples were collected from red grouse killed during legal game bird shoots within the open season in Scotland, from gastrointestinal samples that would normally be discarded as waste material. Therefore, approval for this study from an Animal Welfare and Ethical Review Body was not required.

### Consent for publication

Not applicable

### Availability of data and materials

The datasets generated and analysed during the current study (fastq files, assemblies, genome bins and metagenome assembled genomes) are available in the European Nucleotide Archive under the project number PRJEB85541.

### Authors’ contributions

The authors confirm contribution to the paper as follows: study conception and design - LG; data collection - AA, NH and KF; analysis and interpretation of results - AA and LG; writing – original draft - LG; writing – editing and review - AA, LG, NH and KF. All authors reviewed the results and approved the final version of the manuscript.

## Acknowledgements

We would like to acknowledge the upland estate staff who provided the red grouse samples.

